# Selection, succession and stabilization of soil microbial consortia

**DOI:** 10.1101/533604

**Authors:** Elias K. Zegeye, Colin J. Brislawn, Yuliya Farris, Sarah J. Fansler, Kirsten S. Hofmockel, Janet K. Jansson, Aaron T. Wright, Emily B. Graham, Dan Naylor, Ryan S. McClure, Hans C. Bernstein

## Abstract

Soil microorganisms play fundamental roles in cycling of soil carbon, nitrogen and other nutrients, yet we have a poor understanding of how soil microbiomes are shaped by their nutritional and physical environment. Here we investigated the successional dynamics of a soil microbiome during 21-weeks of enrichment on chitin and its monomer, *N-acetylglucosamine*. We examined succession of the soil communities in a physically heterogeneous soil matrix as well as a highly mixed liquid medium. The guiding hypothesis was that the initial species richness would influence the tendency for the selected consortia to stabilize and maintain relatively constant community structure over time. We also hypothesized that long term, substrate-driven growth would result in consortia with reduced species richness as compared to the parent microbiome and that this process would be deterministic with relatively little variation between replicates. We found that the initial species richness does influence the long-term community stability in both liquid media and soil and that lower initial richness results in a more rapid convergence to stability. Despite use of the same soil inoculum and access to the same major substrate, the resulting community composition differed greatly in soil compared to liquid medium. Hence, distinct selective pressures in soils relative to homogenous liquid media exist and can control community succession dynamics. This difference is likely related to the fact that soil microbiomes are more likely to thrive, with fewer compositional changes, in a soil matrix compared to liquid environments.

**IMPORTANCE:** The soil microbiome carries out important ecosystem functions, but interactions between soil microbial communities have been difficult to study due to the high microbial diversity and complexity of the soil habitat. Here we successfully obtained stable consortia with reduced complexity that contained species found in the original source soil. These consortia and the methods used to obtain them can be a valuable resource for exploration of specific mechanisms underlying soil microbial community ecology. The results of this study also provide new experimental context to better inform how soil microbial communities are shaped by new environments and how a combination of initial taxonomic structure and physical environment influences stability.

## INTRODUCTION

Soil microbiomes are among the most diverse microbial communities on the planet (1, 2) and the majority of soil microbes have not yet been cultivated or studied under laboratory conditions. This, and other confounding properties, such as extreme spatial heterogeneity, make it difficult to study how soil microorganisms interact within natural communities (3). Despite this, a deeper understanding of the ecological properties that control the structure and function of soil microbiomes is needed as they underpin almost every terrestrial food web (4), regulate many elements of Earth’s biogeochemical cycles (5) and are fundamental for growth of healthy crops and bioenergy feedstocks (6).

Well established studies estimate that annual CO_2_ emissions from soil microbial respiration are ten times greater than the CO_2_ produced by fossil fuel utilization (5). Therefore, small changes in the soil carbon cycle – a process linked to microbiome functioning – can have large impacts on atmospheric CO_2_ concentrations. The cycling of complex biopolymers that are both produced and stored in soils largely influences the flux of CO_2_ to the atmosphere. Of these, chitin, an insoluble β-1, 4-linked polymer of *N-acetylglucosamine* (NAG) (7, 8), is a major substrate for soil microbial activity (9) and represents a linkage between the carbon and nitrogen cycles in soils (10-12). Chitin is omnipresent in soil and is an important biopolymer synthesized by fungi (13) and many insects. However, little is known about how chitin and NAG can select for soil-specific bacterial and fungal taxa and influence the structure of microbial communities that are involved in their decomposition.

Successional dynamics of soil microbiomes are related to changes in substrate availability and are crucial to predicting ecosystem development (14–20). During primary succession, early colonizing taxa shape available niche space by regulating pH and nutrient availability (16, 17, 21). However, the feedbacks and processes driving successional patterns constitute fundamental knowledge gaps in understanding trajectories of ecosystem development (16, 19). Microbial succession patterns can be influenced by available resources, including nutrient pools (19, 22), physiochemistry (23), and vegetation (24). Additionally, it is well known that soil moisture is a key determinant of microbial metabolism (25–27). Less is known about how the physical environment, with respect to soil or liquid-like conditions, affect microbial community succession and stability. The relative stability of microbial communities through early succession and thereafter is key to understanding and predicting microbial responses to perturbation (28–31). While the immense complexity of soil microbiomes has hindered many efforts to describe the succession dynamics to ecosystem functioning, organic matter chemistry has been identified as a key driver of primary succession (32).

In this study, we aimed to investigate processes underlying soil microbial community succession by monitoring microbial community development in a sterile soil matrix enriched with NAG. Comparisons were made over the course of 15-weeks of succession to a liquid medium culture derived from the same inoculum. In this way, environmental successional trajectories of the soil microbiome were directly compared to community development using traditional, liquid-based culturing methods that omit the heterogenous chemical and spatial landscapes associated with the soil matrix.

We hypothesized that initial species richness would influence the succession of the consortia and their ability to stabilize with a relatively constant taxonomic structure over time. Specifically, we anticipated that consortia with lower species richness during the initial phases of succession would display higher tendencies to converge towards smaller changes in community structure between successive time points. We also hypothesized that long term selection by NAG would result in soil microbial consortia with reduced complexity as compared to the parent soil microbiome and that this process would be deterministic with relatively little variation between replicates during enrichment.

To test these hypotheses, we investigated the influence of initial richness and physical environment on the progression of chitin/NAG enriched soil microbial consortia. We designed soil microbiome enrichment experiments with the expectation that dilution and long-term selection on chitin/NAG would dramatically reduce the initial community richness when compared to the native soil. One of our aims was to use this procedure to obtain simplified, naturally adapted consortia that can serve as a valuable experimental resource that can be shared for recapitulating some soil microbiome behaviors. We also expected and found that the consortia that did emerge from this long-term succession experiment showed distinct differences based on the physical environment (soil verses liquid). This study has improved our understanding about the succession and stability of microbial communities in soil. Generally, these results show that the final stability of and the extent of species richness were directed by the length of succession, the initial richness, and the culturing environment.

## RESULTS

### Enrichment of a native soil microbiome on chitin

Native soil was supplemented in triplicate with 3 concentrations of chitin (10, 50 and 100 ppm) for six-weeks to select for naturally co-existing soil populations capable of using chitin as a carbon and/or nitrogen substrate. Respiration was monitored during the enrichment as a proxy for soil microbial activity during chitin decomposition. The highest respiration was observed for the highest chitin concentration and therefore the 100 ppm treatments were used to inoculate longer term enrichments supplemented with NAG.

The dominant bacterial phyla in the native soil communities were *Proteobacteria, Actinobacteria, Acidobacteria, Chloroflexi* and *Bacteroidetes*; there were few archaea identified in high relative abundance (Fig. 1) (Supplemental Fig. S1A). The dominant fungi were *Ascomycota* (Supplemental Fig. S1B). The native soil bacterial richness had a mean of 818.5 ± 75.6 16S OTUs (Supplemental Fig. S1C). The native soil bacterial community also exhibited high evenness, with a Simpson’s Evenness score of 0.3; the most abundant OTU accounted for less than 4% of all observations. In comparison to the 16S results, the fungal richness and evenness in the native soil were much lower (Supplemental Fig. S1D). The mean ITS OTU count was 128 ± 39, with a Simpson’s Evenness score of only 0.069, and the top two OTUs together comprised 41% of the observed fungal community.

**Figure 1:**
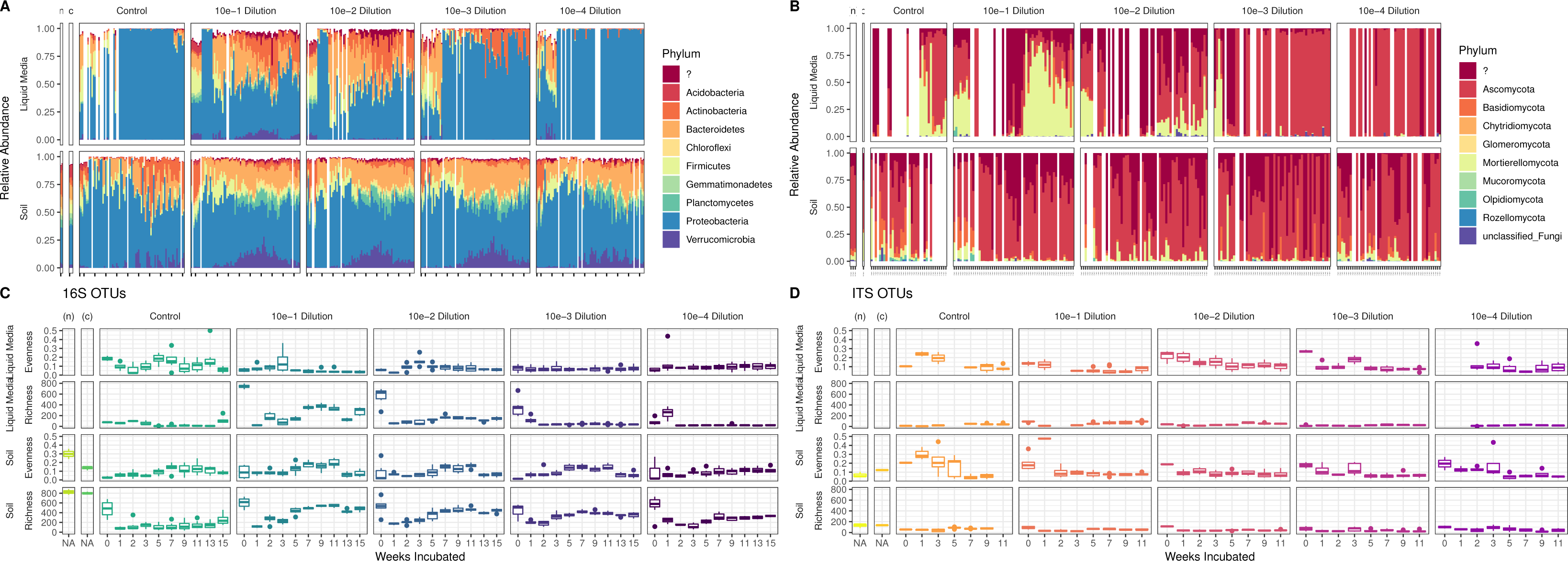
The successional dynamics of microbial consortia. Differences in microbial community structure and alpha diversity are plotted with respect to native soil communities (labelled as ‘n’), the 6-week chitin enriched communities (labelled as ‘c’), uninoculated control (labelled as ‘control’) and different serial dilutions of NAG enrichment in liquid and soil treatments for 0-15 weeks. The results are partitioned by initial dilution and incubating conditions (NAG-enrichment in liquid on the top and soil in the bottom). The most abundant bacterial (A) and fungal (B) phyla are shown over the 15-week incubation. The alpha diversity of bacteria (C) and fungi (D) are estimated using species richness and Simpson’s evenness.

Following the initial 6-week chitin enrichment, the bacterial community structure shifted to a higher relative abundance of *Firmicutes* and *Acidobacteria* and less *Actinobacteria,* although *Proteobacteria* continued to maintain the highest relative abundance (Supplemental Fig. 1A). Additionally, there were shifts in the fungal communities, with a higher relative abundance of *Mortierellomycota* and decreased *Ascomycota* compared to the native soil (Supplemental Fig. 1B). Bacterial richness remained statistically unchanged (p = 0.7624, native soil = 823.5 ± 78.5; n = 2, chitin enriched soil = 801.7 ± 26.35; n = 3). However, bacterial evenness decreased by 55%, indicating that chitin supplementation selected for a subset of populations. The fungal species richness and evenness remained essentially unchanged by the chitin supplementation indicating that the native fungal taxa were less responsive to chitin compared to the bacteria.

### The structure and taxa of soil and liquid-based consortia

After chitin enrichment, subsequent extended enrichment was carried out over 15-weeks using 100 ppm NAG as the major carbon and nitrogen source. The enrichments were performed in two parallel tracks using the same source inoculum (soil enriched for 6-weeks with 100 ppm chitin); in both gamma-irradiated (sterile) soil and liquid M9 medium. The total time for the experiment using chitin enrichment followed by NAG enrichment was 21-weeks. This experimental design was used to optimize opportunities for selection of reduced complexity, naturally co-existing soil consortia and to determine the influence of the physical matrix on the enrichment process. While the physical differences between soil and liquid are paramount, it is important to note other differences including carbon/nitrogen sources or pH that may also have an effect on the succession of resulting consortia.

The NAG enrichments were initiated by serial 10-fold dilution of the chitin enriched soil (dilutions ranged from 10^-1^ to 10^-4^) into the irradiated sterile soil and into liquid M9, both containing 100 ppm of NAG. The relative abundances of both 16S and ITS OTUs differed between serial dilutions and treatment conditions over the course of the experiment (Fig. 1). *Proteobacteria* remained the dominant bacterial phylum during the succession period in both the liquid and soil treatments (Fig. 1A). However, the NAG enriched liquid environment showed a greater degree of change compared to proportions of the most abundant taxa in the native source soil. In the NAG enriched liquid medium, members of the *Proteobacteria* and *Ascomycota* phyla dominated the bacterial and fungal communities, respectively. In contrast, there was a higher diversity of phyla represented in the NAG enriched soil environment over time. In these samples we observed increases in typical soil bacteria that are generally difficult to cultivate; namely *Planctomycetes* and *Verrucomicrobia. Planctomycetes* was negligible in all matching liquid incubations and *Verrucomicrobia* was only present to a comparable degree in the least diluted liquid sample (10^-1^). Simultaneously, we observed depletion of *Acidobacteria* and *Actinobacteria* in the NAG enriched soil. We also detected a greater number of fungal phyla in communities grown on the NAG enriched soil compared to its liquid counterpart, with relatively high proportions of *Mortierellomycota*, *Basidiomycota* and unidentified fungi at the end of the incubation period (Fig. 1B).

As the 15-week NAG enrichments were being regularly sampled for gDNA and respiration, we employed sterile controls to monitor contamination (Supplementary Fig. S2). This enabled us to detect cross contamination between samples as growth in our soil and liquid media controls. This was inferred from non-zero respiration measurements and the recovery of gDNA from liquid media (gDNA was always present in sterile soil). The cross contamination was first observed at week-5 (Supplemental Figure S2). The most common OTU identified from the controls was of the genus *Pseudomonas*. This OTU was present in the native soil and chitin enrichments, indicating that it was intrinsic to the experiential system and native to the parent microbiome (Supplemental Figure S3). Although the sterile controls lacked any viable growth at the onset of the incubations (as determined by plate counting), the *Pseudomonas* OTU introduced during the incubations was able to grow and dominate the liquid sterile controls as well as the more dilute liquid samples (10^-3^ and 10^-4^). However, although present, the *Pseudomonas* OTU did not establish itself to high relative levels within the higher richness liquid samples or any of the soil samples, likely due to the complexity and stability of the existing microbial communities already present in these sites.

We anticipated that long-term selection by NAG in a sterile soil or liquid M9 environment would result in soil microbial consortia with reduced complexity as compared to both the native soil microbiome and the chitin enriched soil microbiome. Over all, this was found to be true although the initial species richness of the inoculum also played a major role. We manually reduced the complexity of the inoculum by controlling the initial species-richness through dilutions. A comparison of the species-richness measured on the first sampling date (week-0) across dilutions in NAG enriched liquid media showed that the dilutions were successful in reducing the richness of the initial inoculum (Fig. 1C and Supplemental Table S1). It is very likely that a corresponding initial drop in richness was also happening with the soil dilutions, although this could not be confirmed by amplicon analysis due to DNA amplification from soil microbes that were likely killed during the gamma irradiation process (33, 34). In the liquid incubations, the observed 16S and ITS OTU counts from the 10^-3^ and 10^-4^ dilutions gradually decreased over time; however, the 10^-1^ and 10^-2^ dilutions revealed sharp decreases in species richness on the first week, followed by a rebounding trend through week-15. This drop and rebounding effect after week-3 was also observed across all of the dilutions associated with the NAG enriched soil. Fungal richness measurements followed similar patterns as those seen for the bacterial richness. By the end of 15-weeks the NAG enriched soil microbiome richness was reduced by approximately 35-70% (depending on dilution) compared to the original native soil (Fig. 1 C, D) and the NAG enriched liquid microbiome richness was reduced by approximately 37-88%. This represents a considerable decrease in species complexity from the initial native and chitin-enriched soil microbiomes and a demonstration that a combination of dilution and long-term selection on specific carbon sources can lead to consortia with reduced species complexity.

### Physical environment and initial species richness influences stability

The stability of the enriched consortia was measured by comparing beta diversity over time. Specifically, we used measures of weighted UniFrac distance (35) between samples that occurred sequentially as a measure of phylogenetic volatility (Fig. 2), where consortia with lower volatility are defined as those showing a more similar community structure from one time point to the next (36). This represents a way to measure how much the community is changing from week-to-week, which is related to the taxonomic compositional stability over time (Supplemental Figure S4). By using this metric, it was clear that while enrichment on both NAG containing soil and liquid media led to stable consortia, those enriched within the liquid environment became relatively stable more quickly as compared to those in the enriched soil (Fig. 2). Consortial stability also depended on the complexity of the initial inoculum (Supplemental Table S2) (Fig. 2), a factor that was controlled by dilution of the chitin enriched input soil. Samples inoculated with the 10^-4^ dilution (lowest initial richness) showed the greatest tendency to stabilize.

**Figure 2:**
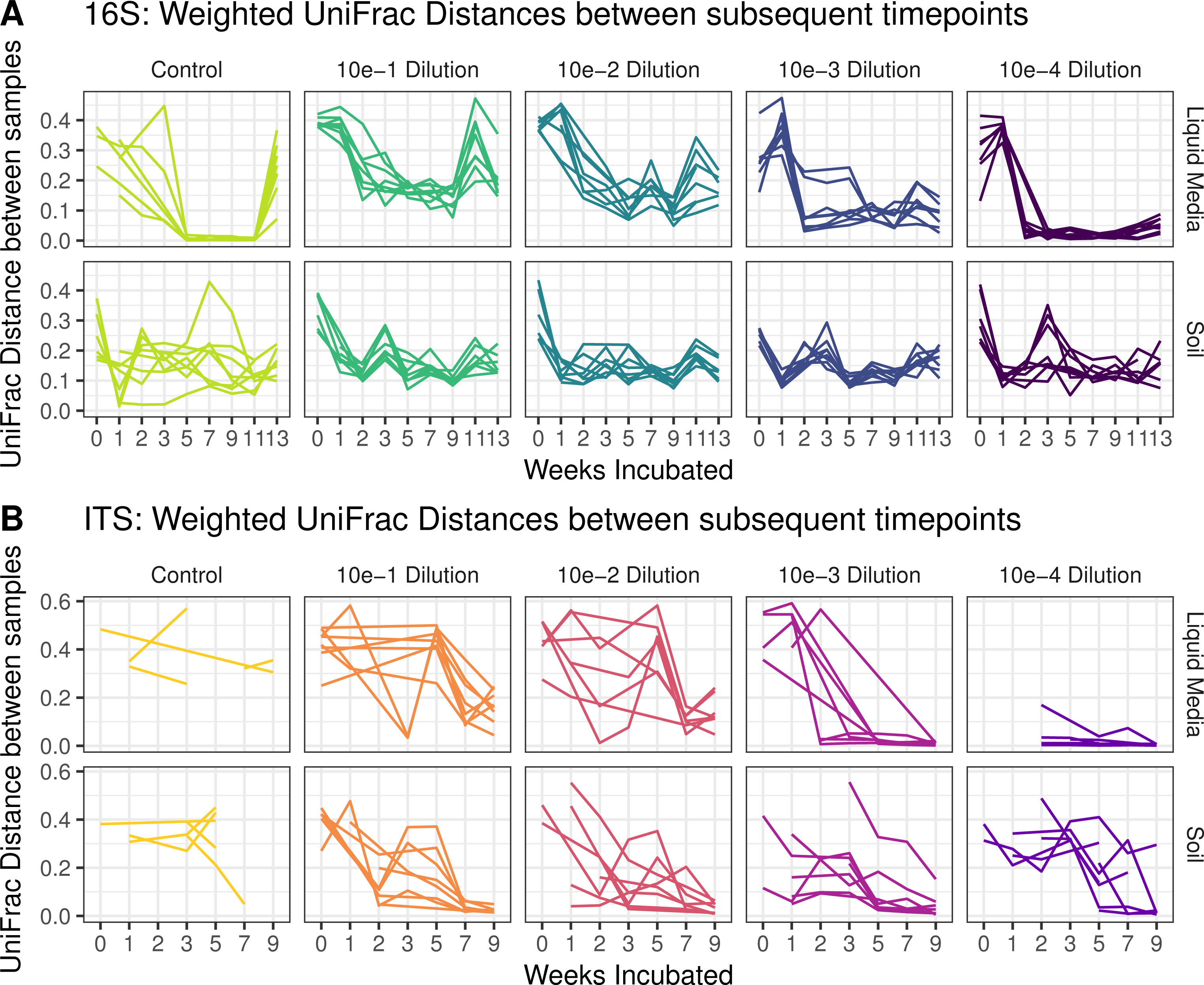
The influence of physical environment on taxonomic volatility – i.e., the tendency for the community to stabilize with respect to taxa being gained/lost over time. Each graph shows the weighted UniFrac distance for bacteria (A) and fungi (B) calculated between subsequent incubation times and plotted by weeks of incubation, dilution factor and treatment conditions.

The consortia became more stable starting at week-5 with the maximum stability reached by the end of the experiment on week-15. However, differences were observed based on succession in liquid versus soil environments. The NAG enriched soil microbial communities showed an initial drop in volatility (weeks 1 – 2) followed by a rise in volatility through weeks 3

– 5 (Fig 2). After 5-weeks of enrichment in soil with NAG, the composition of the soil microbiome did not change significantly (Fig. 2) and volatility continued to drop as the experiment progressed. In contrast, NAG enriched liquid microbiomes initially exhibited an extreme drop in volatility over the first two weeks and thereafter showed either a consistent volatility measurement near 0.10 (dilutions 10^-3^ and 10^-4^) or a continual gradual drop in volatility over the remaining 13-weeks down to a minimum of 0.15 (dilutions 10^-1^ and 10^-2^). Bacterial volatility showed a consistent increase around near week-11, which also corresponded with the observed decreases in the relative abundance of OTUs assigned as *Verruucomicrobia* and *Bacteroidetes* in the soil consortia (Fig. 1). More diverse microbial communities were enriched and stabilized in soil as compared to the liquid incubations. This demonstrates that the physical environment was a significant factor for the stability and compositional convergence of microbial consortia. These results show that the final stability of the consortia and the extent of species richness were directed by the length of succession, initial richness and culturing environment.

### Biological and physical variables underpinning observed beta diversity

Respiration and volatility of the enriched communities were compared to phylogenetic composition over time via ordination by canonical analysis of principal coordinates using weighted Unifrac distance between rarefied samples (Fig. 3). As described earlier, changes (volatility) in community composition between time points were measured as the weighted UniFrac distance between subsequent time points (Fig. 2). In all environments, the volatility vector points were in the direction of early stage incubation samples (Fig. 3), where large changes in the community structure occurred between time points (Supplemental Figure S4). In liquid media incubations, the contribution of respiration for the dissimilarity between samples become more prominent in the later stages of incubation time courses. The dominant phyla from both kingdoms were assessed with respect to incubation time and treatment condition. In the soil, *Proteobacteria* and *Firmicutes* co-varied with volatility (Fig. 3A), as they were most abundant in the volatile initial samples and slowly decreased over time. However, in the liquid media *Firmicutes* and *Bacteroidetes* were closely aligned with volatility (Fig. 3B). Also, *Proteobacteria* became dominant over time in the liquid media incubations; in particular those that were originally inoculated with higher dilutions of the chitin-enrichment that had a lower initial species richness. For the fungi in the soil culture, volatility co-varied primarily with *Mortierellomycota* and *Chytridiomcota*, motile saprotrophs with chitin-containing cells walls that are found in wet soils (Fig. 3C) (37).

**Figure 3:**
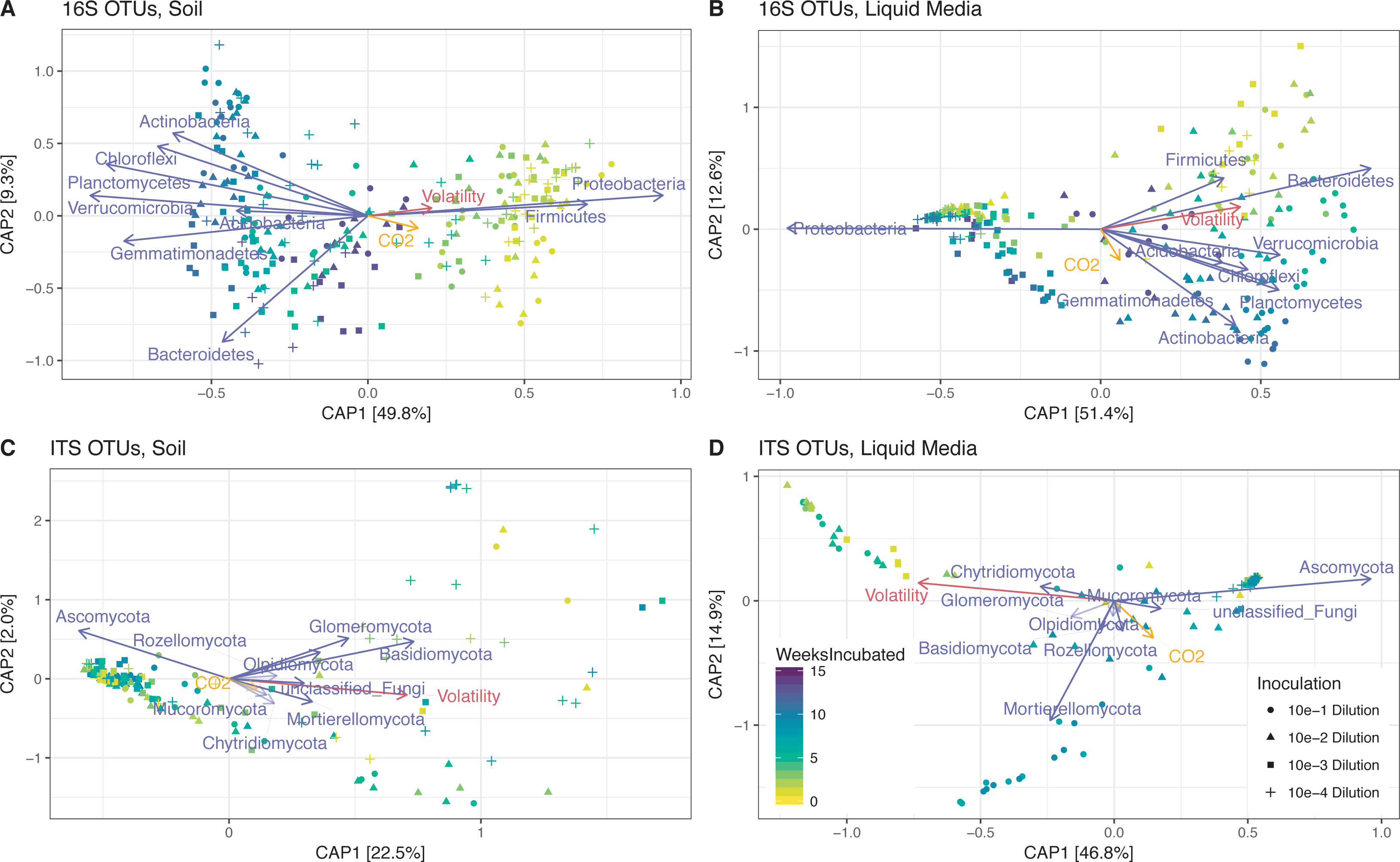
The biological and physical variables correlated with beta diversity. Ordinations present canonical analysis of principal coordinates (CAP) of weighted UniFrac distances between samples. The percent of variation captured by the vectors is shown on each axis. Each vector has a magnitude (length) and direction of a variable's contributions to the principle components. Vectors represent: respiration CO2/h (orange), volatility (red), and the most abundant phyla (purple).

## DISCUSSION

Selective enrichment of soil microbes with specific carbon substrates resulted in the formation of distinct microbial consortia that displayed reduced complexity. Those consortia that developed in NAG enriched soil were also representative of the native soil microbiome used as the inoculum. A primary finding from this study was that the initial species richness influenced successional patterns that were enriched with a specific carbon/nitrogen source in both NAG enriched liquid media and soil incubations. Because the experiment was well replicated (8 biological replicates per treatment) we also confirmed our hypothesis that substrate-driven soil community succession is deterministic in that all of the replicates for a given soil dilution resulted in similar endpoint communities (Supplemental Figure S4 and Table S3). This result was observed in both liquid media and soil substrates, although the taxonomic structure of endpoint consortia was controlled by hydro-physical and other matrix-associated differences between soil and liquid media. The end-point microbial community structure was well explained by the initial dilution condition and this influence was more pronounced on the liquid than the soil treatment condition (Supplemental Table S2). At the end of the enrichment period, the soil NAG enrichments showed higher species richness compared to equivalent liquid treatments, despite having identical inoculations, and were also more representative of microbiomes from the original native soil with respect to community composition. The persistence of members of the original soil microbiome was consistent across dilutions for the NAG enriched soil. The 10^-1^ dilution had the highest species similarity to the native soil and retained a diverse community, while, at the other extreme, the 10^-4^ dilution represented a much simpler, less rich community.

This approach for developing consortia with reduced complexity is of interest as a method for obtaining simplified model microbiomes, with naturally interacting members that are representative of the native soil system. This similarity to the native soil seen with the NAG enriched soil is likely a result of the experiment being performed with soil microbes in their natural soil substrate, compared to a relatively foreign substrate (liquid). Another recent study also examined shorter term succession of soil microbiomes in liquid (but not soil) and found that soil microbiomes enriched on liquid media are very different from the original source soil microbiome (38). That study was carried out using a variety of carbon sources resulting in microbiomes with reduced complexity, similar to what we show here. Together, these studies confirm that reduced complexity consortia that have community membership representative of soil microbiomes are much more likely to be obtained using a soil-based enrichment compared to a liquid-based enrichment. In addition, our results clearly show differences in the successional dynamics and end-point structures of each consortium with respect to their initial species richness based on the dilution of the chitin enriched soil inoculum.

We found that the richness of the initial soil inoculum strongly impacted the alpha diversity of the resulting microbial consortia over time (Fig. 1). These results support our hypothesis that the initial species richness would influence each consortium’s tendency to converge towards smaller changes in community structure between successive time points. Results supporting this hypothesis were observed for the higher dilutions (10^-3^ and 10^-4^) for all treatments and measurements (Fig. 2). Each consortium’s tendency to converge towards smaller changes in community structure between successive time points was assessed by comparing weighted UniFrac distances between time points and was notably strongest for communities developed in the liquid media and measured by 16S rRNA sequencing as compared to ITS. The generalizability of this stability convergence effect is partially supported through similar findings presented by Shade et al (2014), who showed how rare taxa significantly influence microbial diversity (39). Dilutions are more likely to remove rare taxa and therefore our results provide some additional quantification of the effect presented by Shade et al. (2014). However, in our current study, we cannot fully decouple the effects of reduced initial richness from reduced counts of viable cells that were almost certainly created from the dilution procedure. Hence, an alternative interpretation could be formulated as decreased viable cell numbers in early stages of succession lead to decreased species richness and higher tendencies to converge towards smaller changes in community structure between successive time points.

Both bacterial and fungal populations were selected during the chitin/NAG incubation process. This suggests that the representative populations were able to either metabolize, or through some other means take advantage of the added substrates. Specifically, we found that members of the *Acidobacteria, Actinobacteria, Bacteroidetes, Chloroflexi, Firmicutes, Gemmatimonadetes, Planctomycetes, Proteobacteria, Verrucomicrobia* were represented in the NAG incubations (Fig. 1a). In addition, the richness of *Verrucomicrobia*, *Bacteroidetes* and *Planctomycetes* increased in soil incubated with NAG compared to native soil (Fig. 1a). Representatives of these phyla were also detected on a previous study of soil enriched with chitin (12). With respect to the fungi, we found that the *Mortierellomycota* phylum increased in relative abundance in the NAG enriched consortia (Fig. 1b). *Mortierellomycota* are members of the Mucoromyceta, based on recent fungal taxonomy (40). They are sporangiferous, generally saprotrophic, including being able to grow on other fungi, and are found in soil (41). The dominance of these specific bacteria and fungi suggests that their enrichment came due to their ability to use either chitin/NAG or its metabolic byproduct as a substrate.

The occurrence of enriched, stable consortia with dozens to hundreds of members, found here and in a similar study by Goldfarb, et. al. (42), as opposed to selection of a monoculture, suggests that the compositions of the reduced microbial communities are governed by cross-feeding interactions among microbes. In our longer-term soil incubations with NAG the microbiome converged into a less complex microbial community compared to that found in the native soil. This is consistent with the results of the Goldfarb study, which also enriched a simplified microbial community, derived from soil, on single carbon sources. However, unlike the previous study, which used only liquid, we enriched on both liquid and soil and found that enrichments on soil led to a reduced complexity community that is far more representative of the native soil microbiome compared to enrichment on liquid. There are several reasons why structured environments may better facilitate and stabilize social interactions, including the limited dispersal of interacting species, and the physical retention of resources within the soil matrix. The close physical proximity of members of soil consortia in discrete niches would thus facilitate social activities between member populations (e.g. exchange of public goods, quorum sensing and competition). When microbial communities have a single major carbon source only a subset of the community will have the metabolic capability to utilize it as a substrate. For complex substrates, such as chitin, other species will be reliant on primary species to degrade the polymer to simpler compounds, thus selecting for a community that interacts by metabolic cross-feeding, interactions that positively affect both the primary degrading species and the secondary degrading species (43, 44). Positive metabolic interactions between microorganisms residing within communities have been studied in other systems as well; particularly in biofilms where species and cells are in very close proximity and must cooperate for growth (45).

Because we monitored the soil enrichments over a relatively long time period, we could determine the time required for the soil microbiomes to reach stable community memberships. Stability was achieved surprisingly rapidly (3 – 5 weeks) and the resulting consortia remained stable over several months. Importantly, the development of stable, reduced complexity, naturally interacting consortia from native soil can provide representative model soil communities for future studies to study the mechanisms underlying species interactions. This valuable resource should enable deciphering of the molecular signaling mechanisms and metabolic interactions used by soil community members to decompose complex carbon substrates in soil. In addition, the information can be used to enhance *in silico* models of soil microbial community interactions that can be used to predict how key taxa and traits can be perturbed by environmental change.

### Conclusions

Here we demonstrated that the succession of microbial communities derived from chitin/NAG enriched soil microbiome is strongly influenced by the initial soil microbiome richness and the hydro-physical environment. The initial species richness, which is a proxy for the complexity of a microbiome, at least partially controlled the tendency for a soil-sourced consortium to stabilize and maintain a relatively constant community structure over time. Additionally, the long-term soil enrichments resulted in a reduced complexity representation of the initial soil microbiome diversity and richness. The results of this study inform how soil microbial communities are shaped during succession and how a combination of the initial taxonomic structure and physical environment influences the tendency for a community to stabilize over time.

## MATERIALS AND METHODS

### Field sampling and chitin enrichment

Soil was collected in October 2017 from a field site operated by Washington State University, located in Prosser, Washington State USA (46°15’04”N and 119°43’ 43”W) site. The soil represents a Warden silt loam that is characterized as a coarse-silty, mixed, superactive and mesic Xeric Haplocambid. The soil represents a marginal soil with low organic matter content (3.7%) and pH = 8. All soil samples were collected in three field replicates. At each site, bulk sampling was accomplished with a shovel within 0 to 20 cm depth from the ground and it was stored in plastic bags at 4°C. To exclude bigger soil aggregates and rocks, samples were sieved with 4 mm mesh size. For each of the three field blocks, three homogeneous replicates of 150 g soil were weighed out in to a 250 ml sterile screwcap bottles. To enrich chitin degrading members of the microbial community, samples were incubated for six weeks in soil augmented with chitin (*Poly-(1→4)-β-N-acetyl-D-glucosamine,* Sigma-Aldrich, St. Louis, MO, USA) at different concentrations (0, 10, 50,100 μg chitin/g soil dry weight). Chitin was mixed and evenly distributed within the soil, and sterile water was added to reflect a 24% field water capacity. Samples were kept in the dark at 20°C. Additionally, 1 g of sample was harvested weekly from each bottle and stored in -80°C for 16S and ITS amplicon analysis.

### Gamma irradiated soil

Prosser soil was sterilized with gamma irradiation at 85 kGy additional in two successive applications of or 25 kGy, followed by 60 kGy. Initially, 3000 curie Co-60 source was used in the collimated open beam irradiator. For the second irradiation, 1300 curie Co-60 source was used in the Gamma Bunker, which is a 1.5 ft.^3^ closed chamber irradiator (46, 47). Sterility of soil was confirmed by plating of several serial dilutions on LB agar plates followed by incubation at 30 °C and the lack of growth.

### Sterile soil incubations and liquid controls

M9 minimal media and sterile liquid soil extract were prepared as described by Sambrook and Russell (2001) (48) and method of soil analysis-microbiological and biochemical properties (49), respectively. *N-acetyl glucosamine* (NAG) (Sigma-Aldrich, St. Louis, MO, USA) was added into the M9 media to 100 μg/ml. Ten milliliter liquid cultures were setup in 25 ml sterile glass tubes in four successive 10-fold serial dilutions. First, 1 g of actively respiring chitinolytic enriched soil (100 μg chitin/g soil dry weight) was inoculated into the first glass tubes with 9 mL of the M9 media (representing the 10^-1^ dilution) and vortexed for 30 seconds. This solution then was used for the subsequent serial dilutions. Uninoculated controls were also generated and incubated with the dilution samples. Each serial dilution and respective controls were performed in 8 biological replicates. Incubation was performed in the dark at 20 °C, shaking at 130 RPM. CO_2_ respiration was measured aseptically three times a week. Headspace was aseptically flushed with air after each sample to prevent anaerobic conditions. Additionally, 1 ml of sample was harvested weekly for the first three weeks, followed by biweekly sampling and stored in -80 °C for 16S and ITS amplicon analysis. After each sampling period, substrate and moisture levels were refreshed by adding 1 ml of M9 medium.

The soil enrichments were set up using 5.5 g gamma-irradiated soil in 15ml sterile tubes in parallel with their liquid counterparts. The “sterile soil” treatments were prepared by 1 ml soil extract liquid enriched with 100 ppm NAG to each tube containing sterile soil. The soil samples were briefly mixed with a sterile spatula and pre-incubated in the dark at 20°C for two days. After pre-incubation, 0.5 ml inoculum was taken from the liquid serial dilution described above and added to the counterpart sterile soil tubes. The soil enrichments were sealed with filter screw caps (CellTreat, non-pyrogenic and sterilized by gamma irradiation, China) to allow continuous air flow. Each 0.3 g sample was harvested weekly for the first three weeks, followed by biweekly sampling and stored at -80 °C for downstream molecular measurements.

### Amplicon sequencing

Total DNA was extracted using the MoBio PowerSoil DNA isolation kit (Qiagen, Carlsbad, CA) in accordance with the Earth Microbiome Project (EMP) protocols (50). Sequencing was performed on an Illumina MiSeq instrument (Illumina, San Diego, CA). Triplicate, separate 16S and ITS rRNA gene amplification reactions were performed on DNA from each extraction. The 16S primers targeted the V4 hypervariable region of the 16S SSU rRNA gene using the V4 forward primer (515F) and V4 reverse primer (806R) with 0–3 random bases and the Illumina sequencing primer binding site (51). The ITS primers targeted the ITS1 region using the ITS1f and ITS2 primers (52).

### Amplicon analysis

The Hundo (2017) amplicon processing protocol was used to process 16S and ITS amplicons (53). In brief, sequences were trimmed and filtered of adapters and contaminants using BBDuk2 of the BBTools (‘Tools’) package. VSEARCH (54) was used to merge, filter to an expected error rate of 1, dereplicate, and remove singletons before preclustering reads for *de novo* and reference-based chimera checking. Reads were clustered into OTUs at 97% similarity and an OTU table in the BIOM format (55) was constructed by mapping filtered reads back to these clusters. BLAST+ (56) is used to align OTU sequences to the database curated by CREST (57) (SILVA v128 for 16S, UNITE v7 for ITS) and taxonomy was assigned based on the CREST LCA method. Multiple sequence alignment was performed with Clustal Omega (58) and a phylogenetic tree was constructed using FastTree2 (59).

### Diversity analysis

Downstream analysis was completed in R (60), using the phyloseq (61) and vegan packages (62). To preserve the maximum consistency within each replicate, samples were rarified to an even depth of 2,000 reads per sample. The observed counts of unique OTUs (species-richness) and Simpson’s Evenness were used to characterize alpha diversity (63). In order to assess microbial stability/volatility over time, we implemented the volatility analysis as previously described (36), in which the amount the community change between successive time points was measured with weighted UniFrac distances.

### Data repository and reproducible analyses

Genetic sequencing data is available on the Open Science Framework (osf.io) for both 16S and 1TS amplicons as part of this project: https://osf.io/6d5kz/, along with the Rmarkdown processing scripts used to process the data and build annotated feature abundance tables and phylogenic tree using Hundo.

## ACKNOWLEDGMENT

This research was supported by the Department of Energy Office of Biological and Environmental Research (BER) and is a contribution of the Scientific Focus Area ‘Phenotypic response of the soil microbiome to environmental perturbations’ (70880). PNNL is operated for the DOE by Battelle Memorial Institute under Contract DE-AC05-76RLO1830.

---

**Supplemental Figure S1:**
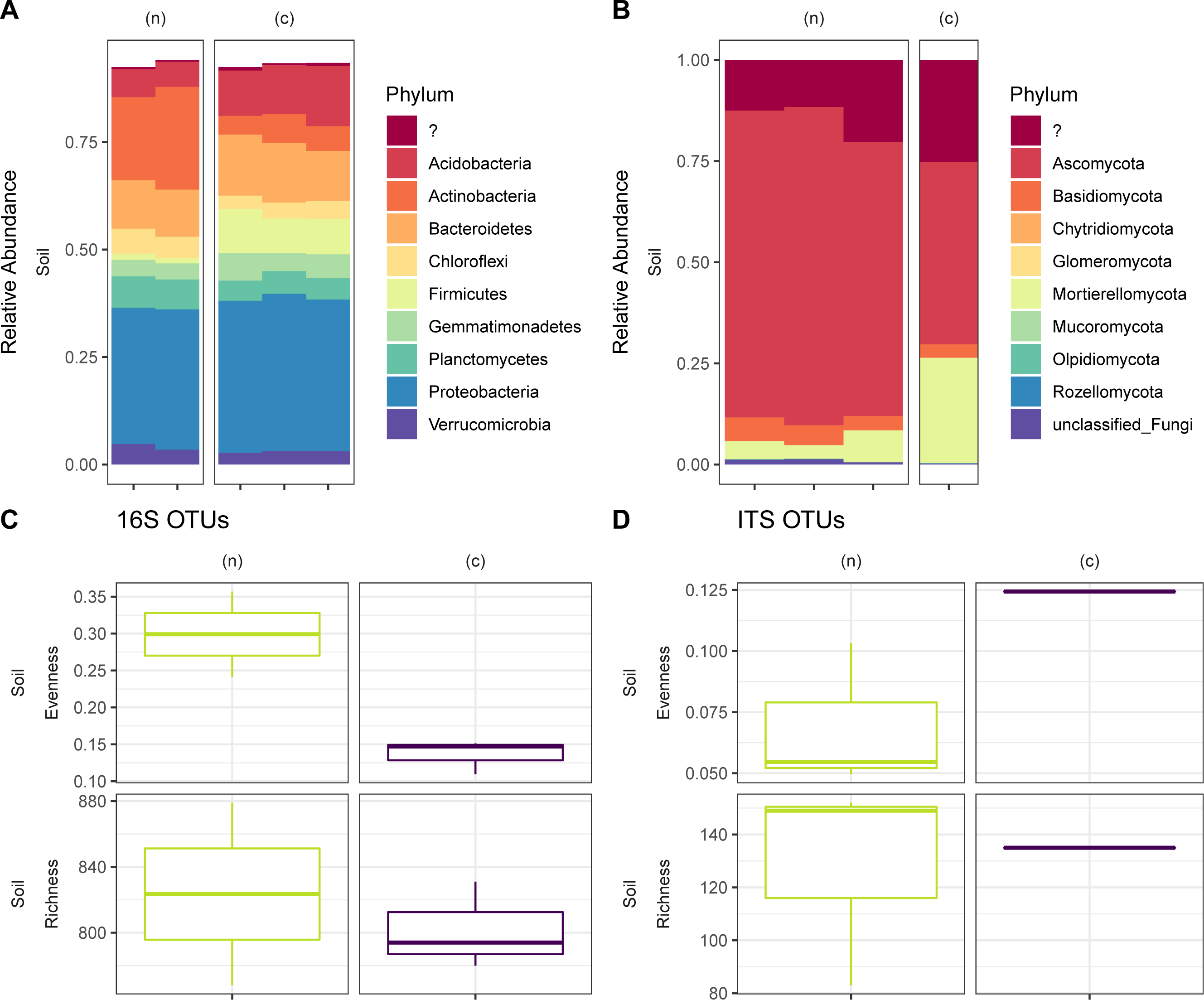
The most abundant bacterial (A) and fungal (B) phyla are plotted for the native soil (labelled as ‘n’) and the 6-week chitin enriched soil (labelled as ‘c’). The alpha diversity of bacteria (C) and fungi (D) are estimated using species richness and Simpson’s evenness.

**Supplemental Figure S2:**
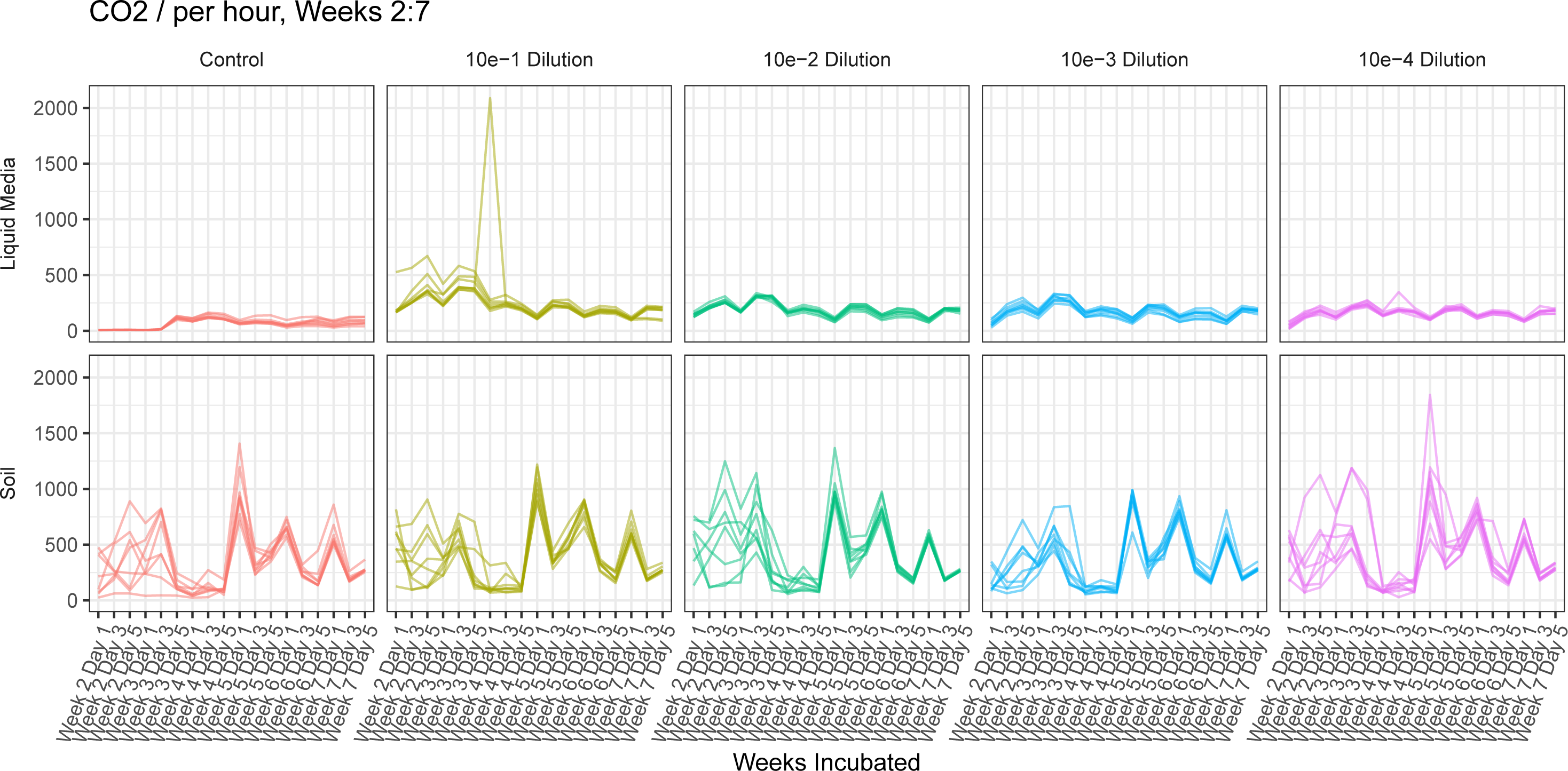
Rate of respiration varies over time based on dilution and physical environment. The respiration rate was plotted for both liquid (top) and soil (bottom) NAG enrichment cultures from weeks 2 through 7 to avoid the light spike in CO_2_ released immediately after inoculation. Respiration was monitored three times a week with a PP Systems-EGM-4 Environmental Gas Monitor (Amesbury, MA). The gas analyzer was setup in static mode to measure CO_2_ and monitor respiration in each sample at periodic intervals. To measure CO_2,_ a 10 mL gas aliquot was taken from the incubation container using a sterile plastic syringe which has a control valve.

**Supplemental Figure S3:**
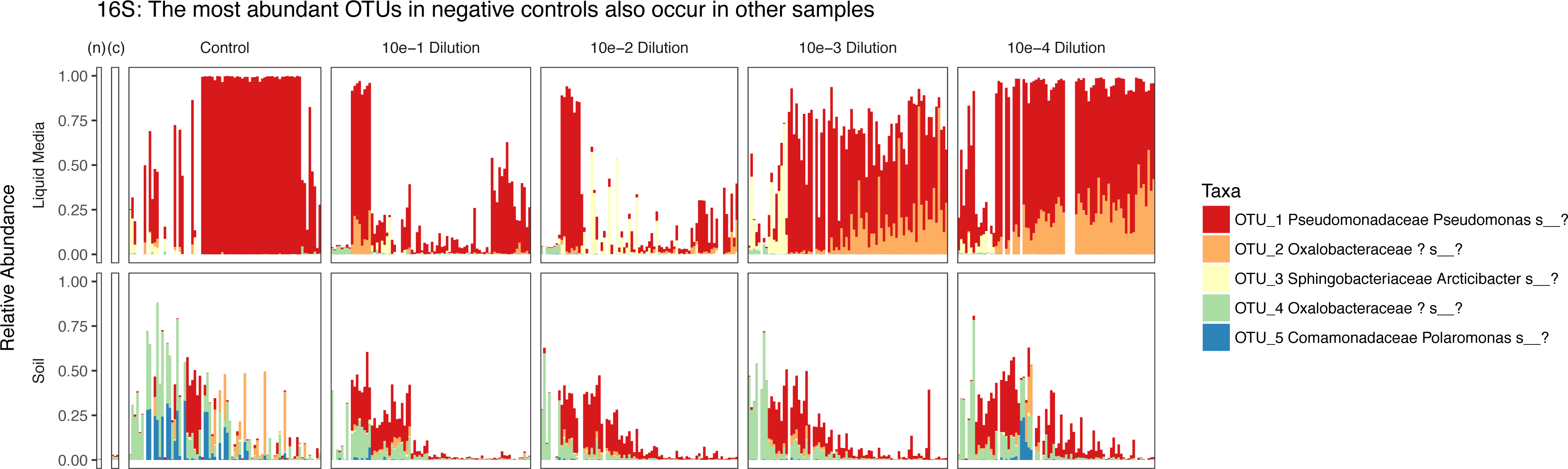
Bacterial OTUs observed from 16S rRNA gene amplicon products in uninoculated controls. The most abundant OTUs shown in the controls were also found in the other samples.

**Supplemental Figure S4:** Patterns of variation within sample groups and within replicates. A) Beta dispersion (within sample group variation) varies among time points as shown by the pairwise permutation test for homogeneity of multivariate dispersion at each time. Pairwise comparisons of the observed p-values are tabulated for both the 16S and ITS for liquid and soil treatment conditions. B) A linear regression fit to the pairwise distance between replicates also shows changes in dispersion over time.

**Supplemental Table S1:** Final richness significantly varies between dilutions. Tukey’s HSD test was performed to compare the richness between dilutions, and the p-values are shown here.

**Supplemental Table S2:** An initial time point species-richness in NAG enriched liquid media across all dilutions and control.

**Supplemental Table S3:** Community composition of final timepoints can be explained by initial dilution. On each set of endpoints, the adonis test was used to measure how much variation in weighted UniFrac distances between samples could be explained by the initial treatment (R^2^) and the probability that this level of similarity between endpoints would be seen by chance alone (p-value).

**Table S1.** Final richness significantly varies between dilutions. Tukey's HSD test was performed to compare the richness between dilutions, and the p-values are shown here.

| Type | Treatment | 10e-2 Dilution -<br>10e-1 Dilution | 10e-3 Dilution -<br>10e-1 Dilution | 10e-3 Dilution -<br>10e-2 Dilution | 10e-4 Dilution -<br>10e-1 Dilution | 10e-4 Dilution -<br>10e-2 Dilution | 10e-4 Dilution -<br>10e-3 Dilution |
| --- | --- | --- | --- | --- | --- | --- | --- |
| 16S | Soil | 0.002598 | 0 | 0 | 0 | 0 | 0.1138814 |
| 16S | Liquid | 0 | 0 | 0 | 0 | 0 | 0.0000024 |
| ITS | Soil | 0.0007323 | 0 | 0.0000002 | 0.0009251 | 0.9998387 | 0.0000002 |
| ITS | Liquid | 0 | 0 | 0 | 0 | 0 | 0.0014198 |

**Table S2.** Community composition of final timepoints can be explained by initial dilution. On each set of endpoints, the adonis test was used to measure how much variation in weighted UniFrac distances between samples could be explained by the initial treatment (R^2^) and the probability that this level of similarity between endpoints would be seen by chance alone (p-value).

| Type | Treatment | R2 | p.value |
| --- | --- | --- | --- |
| 16S | Inoculated Soil | 0.3109907 | 0.001 |
| 16S | Inoculated Liquid | 0.7656066 | 0.001 |
| ITS | Inoculated Soil | 0.2955797 | 0.008 |
| ITS | Inoculated Liquid | 0.6843260 | 0.001 |

**Table S3.**
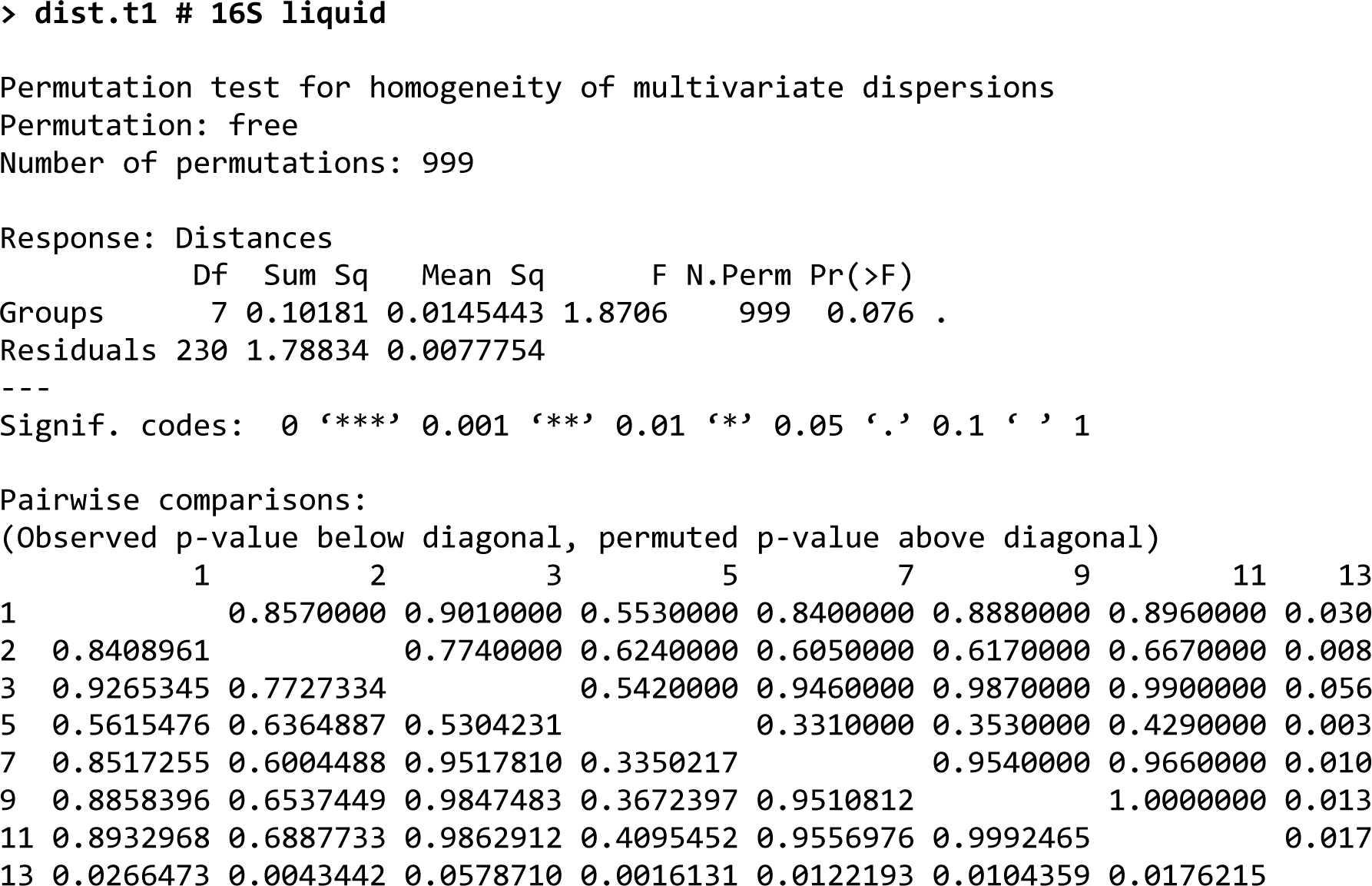
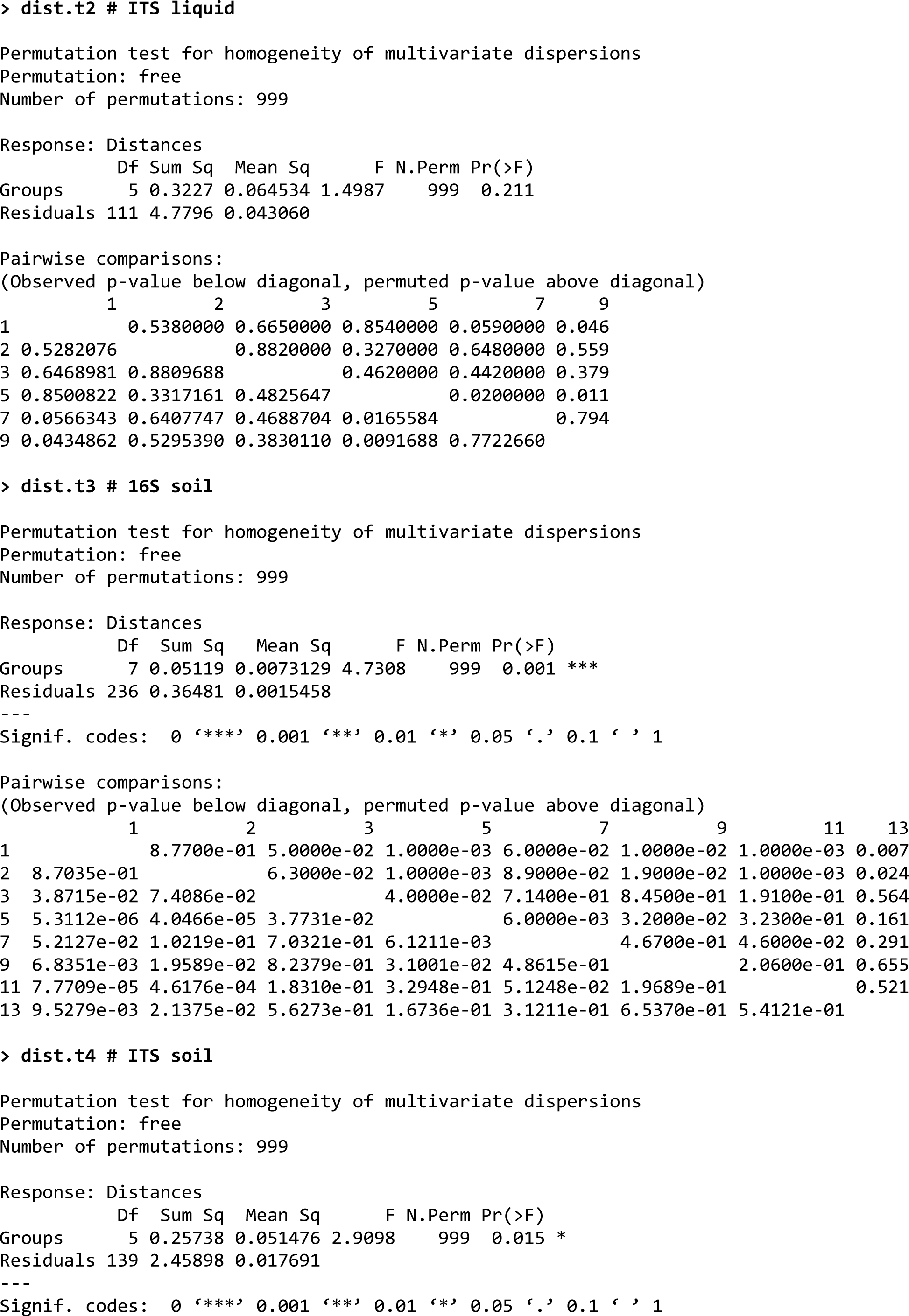
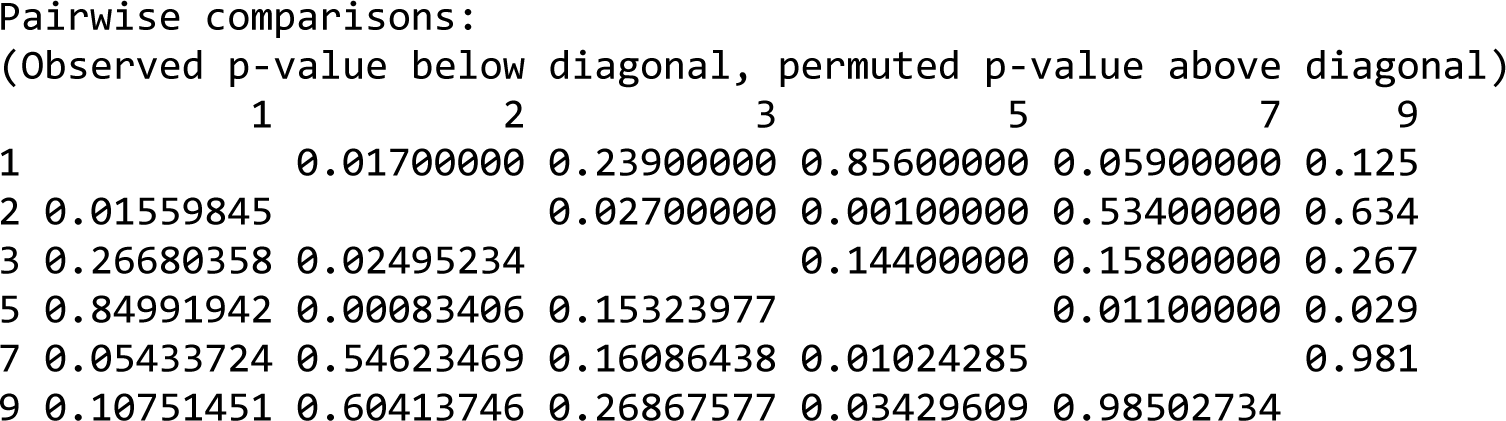
Outputs from the tests ofr homogeneity of multivariate dispersions performed with respect to each treatment.

|  | Df | Sum Sq | Mean Sq | F | N.Perm | Pr(>F) |
| --- | --- | --- | --- | --- | --- | --- |
| Groups | 7 | 0.10181 | 0.0145443 | 1.8706 | 999 | 0.076 |
| Residuals | 230 | 1.78834 | 0.0077754 |  |  |  |

|  | 1 | 2 | 3 | 5 | 7 | 9 | 11 | 13 |
| --- | --- | --- | --- | --- | --- | --- | --- | --- |
| 1 |  | 0.8570000 | 0.9010000 | 0.5530000 | 0.8400000 | 0.8880000 | 0.8960000 | 0.030 |
| 2 | 0.8408961 |  | 0.7740000 | 0.6240000 | 0.6050000 | 0.6170000 | 0.6670000 | 0.008 |
| 3 | 0.9265345 | 0.7727334 |  | 0.5420000 | 0.9460000 | 0.9870000 | 0.9900000 | 0.056 |
| 5 | 0.5615476 | 0.6364887 | 0.5304231 |  | 0.3310000 | 0.3530000 | 0.4290000 | 0.003 |
| 7 | 0.8517255 | 0.6004488 | 0.9517810 | 0.3350217 |  | 0.9540000 | 0.9660000 | 0.010 |
| 9 | 0.8858396 | 0.6537449 | 0.9847483 | 0.3672397 | 0.9510812 |  | 1.0000000 | 0.013 |
| 11 | 0.8932968 | 0.6887733 | 0.9862912 | 0.4095452 | 0.9556976 | 0.9992465 |  | 0.017 |
| 13 | 0.0266473 | 0.0043442 | 0.0578710 | 0.0016131 | 0.0122193 | 0.0104359 | 0.0176215 |  |
| Groups | 5 | 0.3227 | 0.064534 | 1.4987 | 999 | 0.211 |
| Residuals | 111 | 4.7796 | 0.043060 |  |  |  |

|  | 1 | 2 | 3 | 5 | 7 | 9 |
| --- | --- | --- | --- | --- | --- | --- |
| 1 |  | 0.5380000 | 0.6650000 | 0.8540000 | 0.0590000 | 0.046 |
| 2 | 0.5282076 |  | 0.8820000 | 0.3270000 | 0.6480000 | 0.559 |
| 3 | 0.6468981 | 0.8809688 |  | 0.4620000 | 0.4420000 | 0.379 |
| 5 | 0.8500822 | 0.3317161 | 0.4825647 |  | 0.0200000 | 0.011 |
| 7 | 0.0566343 | 0.6407747 | 0.4688704 | 0.0165584 |  | 0.794 |
| 9 | 0.0434862 | 0.5295390 | 0.3830110 | 0.0091688 | 0.7722660 |  |

|  | Df | Sum Sq | Mean Sq | F | N.Perm | Pr(>F) |
| --- | --- | --- | --- | --- | --- | --- |
| Groups | 7 | 0.05119 | 0.0073129 | 4.7308 | 999 | 0.001 *** |
| Residuals | 236 | 0.36481 | 0.0015458 |  |  |  |

|  | 1 | 2 | 3 | 5 | 7 | 9 | 11 | 13 |
| --- | --- | --- | --- | --- | --- | --- | --- | --- |
| 1 |  | 8.7700e-01 | 5.0000e-02 | 1.0000e-03 | 6.0000e-02 | 1.0000e-02 | 1.0000e-03 | 0.007 |
| 2 | 8.7035e-01 |  | 6.3000e-02 | 1.0000e-03 | 8.9000e-02 | 1.9000e-02 | 1.0000e-03 | 0.024 |
| 3 | 3.8715e-02 | 7.4086e-02 |  | 4.0000e-02 | 7.1400e-01 | 8.4500e-01 | 1.9100e-01 | 0.564 |
| 5 | 5.3112e-06 | 4.0466e-05 | 3.7731e-02 |  | 6.0000e-03 | 3.2000e-02 | 3.2300e-01 | 0.161 |
| 7 | 5.2127e-02 | 1.0219e-01 | 7.0321e-01 | 6.1211e-03 |  | 4.6700e-01 | 4.6000e-02 | 0.291 |
| 9 | 6.8351e-03 | 1.9589e-02 | 8.2379e-01 | 3.1001e-02 | 4.8615e-01 |  | 2.0600e-01 | 0.655 |
| 11 | 7.7709e-05 | 4.6176e-04 | 1.8310e-01 | 3.2948e-01 | 5.1248e-02 | 1.9689e-01 |  | 0.521 |
| 13 | 9.5279e-03 | 2.1375e-02 | 5.6273e-01 | 1.6736e-01 | 3.1211e-01 | 6.5370e-01 | 5.4121e-01 |  |

|  | Df | Sum Sq | Mean Sq | F | N.Perm | Pr(>F) |
| --- | --- | --- | --- | --- | --- | --- |
| Groups | 5 | 0.25738 | 0.051476 | 2.9098 | 999 | 0.015 * |
| Residuals | 139 | 2.45898 | 0.017691 |  |  |  |

|  | 1 | 2 | 3 | 5 | 7 | 9 |
| --- | --- | --- | --- | --- | --- | --- |
| 1 |  | 0.01700000 | 0.23900000 | 0.85600000 | 0.05900000 | 0.125 |
| 2 | 0.01559845 |  | 0.02700000 | 0.00100000 | 0.53400000 | 0.634 |
| 3 | 0.26680358 | 0.02495234 |  | 0.14400000 | 0.15800000 | 0.267 |
| 5 | 0.84991942 | 0.00083406 | 0.15323977 |  | 0.01100000 | 0.029 |
| 7 | 0.05433724 | 0.54623469 | 0.16086438 | 0.01024285 |  | 0.981 |
| 9 | 0.10751451 | 0.60413746 | 0.26867577 | 0.03429609 | 0.98502734 |  |

